# Optogenetic inhibition reveals distinct roles for basolateral amygdala activity at discrete timepoints during risky decision making

**DOI:** 10.1101/180885

**Authors:** Caitlin A. Orsini, Caesar M. Hernandez, Sarthak Singhal, Kyle B. Kelly, Charles J. Frazier, Jennifer L. Bizon, Barry Setlow

## Abstract

Decision making is a multifaceted process, consisting of several distinct phases that likely require different cognitive operations. Previous work showed that the basolateral amygdala (BLA) is a critical substrate for decision making involving risk of punishment; however, it is unclear how the BLA is recruited at different stages of the decision process. To this end, the current study used optogenetics to inhibit the BLA during specific task phases in a model of risky decision making (Risky Decision-making Task; RDT) in which rats choose between a small, “safe” reward and a large reward accompanied by varying probabilities of footshock punishment. Rats received intra-BLA microinjections of viral vectors carrying either halorhodopsin (eNpHR3.0-mCherry) or mCherry alone (control) followed by optic fiber implants and were trained in the RDT. Laser stimulation during the task occurred during either intertrial interval, deliberation, or reward outcome phases, the latter of which was further divided into the three possible outcomes (small, safe; large, unpunished; large, punished). Inhibition of the BLA selectively during the deliberation phase decreased choice of the large, risky outcome (decreased risky choice). In contrast, BLA inhibition selectively during delivery of the large, punished outcome increased risky choice. Inhibition had no effect during the other phases, nor did it affect performance in control rats. Collectively, these data indicate that the BLA can either inhibit or promote choice of risky options, depending on the phase of the decision process in which it is active.

**Significance Statement:** To date, most behavioral neuroscience research on neural mechanisms of decision making has employed techniques that preclude assessment of distinct phases of the decision process. Here we show that optogenetic inhibition of the basolateral amygdala (BLA) has opposite effects on choice behavior in a rat model of risky decision making depending on the phase in which inhibition occurs. BLA inhibition during a period of deliberation between small, safe and large, risky outcomes decreased risky choice. In contrast, BLA inhibition during receipt of the large, punished outcome increased risky choice. These findings highlight the importance of temporally targeted approaches to understand neural substrates underlying complex cognitive processes. More importantly, they reveal novel information about dynamic BLA modulation of risky choice.

## Introduction

The ability to make adaptive choices requires multiple cognitive operations that work in concert to guide efficient and optimal behavior (Rangel et al., 2008). For example, an organism must calculate the objective and subjective value of the available options, which entails evaluating the relative value of both the benefits and costs associated with each option. This information must be acquired from past experience, such as the contingencies of previous actions and their outcomes, as well as other motivational (e.g., hunger) and environmental (e.g. presence of salient predictive cues) factors. Finally, the organism must determine the value of the actual outcome of its choice, and use this information as feedback to guide future choices. Together, these processes allow an organism to execute or inhibit its choice behavior as appropriate to its past, current, and anticipated future conditions. While the majority of individuals are able to effectively engage these processes and make adaptive decisions, individuals with psychiatric diseases such as substance use disorder, anorexia nervosa, and post-traumatic stress disorder exhibit impaired decision making (Bechara and Damasio, 2002; Crowley et al., 2010; Najavits et al., 2011; Schneider et al., 2012; Kaye et al., 2013; Gonzalez et al., 2015; Dekkers et al., 2016), often resulting in maladaptive choices. The specific components of the decision making process that are perturbed in these pathological conditions, however, are unclear.

Decision making is mediated by interconnected brain structures within the mesocorticolimbic circuit (Orsini et al., 2015b). One such structure within this circuit that has received considerable attention in cost/benefit decision making is the basolateral amygdala (BLA) (Winstanley and Floresco, 2016). Using well-validated rodent models of risk-based decision making, previous work has shown that lesions or pharmacological inactivation of the BLA result in disadvantageous choices (Winstanley et al., 2004; Ghods-Sharifi et al., 2009; Zeeb and Winstanley, 2011; Hosking et al., 2014; Tremblay et al., 2014). This is consistent with neuroimaging data showing that the amygdala is activated during assessment of risky choices (De Martino et al., 2006; Roiser et al., 2009) and is hypoactive in individuals with impaired risky decision making (Crowley et al., 2010; Gowin et al., 2013). More recently, the BLA has been implicated in decision making involving risk of explicit punishment (Orsini et al., 2015a). In well-trained rats, BLA lesions increased choice of a large reward associated with risk of footshock punishment. These data suggested that the BLA is critical for the integration of reward- and punishment-related information to guide optimal behavior. Importantly, however, it is unclear how and at what point in the decision-making process this integration occurs.

*In vivo* electrophysiological studies show that BLA neurons do not respond uniformly to salient stimuli but instead mediate different aspects of motivated behavior. For example, different populations of BLA neurons respond differentially to rewarding and aversive outcomes (Schoenbaum et al., 1998, 1999; Paton et al., 2006; Belova et al., 2007; Belova et al., 2008; Shabel and Janak, 2009; Sangha et al., 2013; Gore et al., 2015), and are organized into intrinsically (Zhang et al., 2013) and extrinsically distinct circuits (Namburi et al., 2015; Beyeler et al., 2016). In addition, amygdala neurons differentially contribute to generation of prospective plans to obtain immediate rewards (Grabenhorst et al., 2012) as well as rewards in the distant future (Hernadi et al., 2015). This functional heterogeneity within the BLA supports the hypothesis that the BLA is differentially engaged during decision making involving rewarding and aversive outcomes. How the BLA is recruited, however, may depend on the specific cognitive components of the decision-making process. In other words, how the BLA contributes to the deliberative process of decision making may be distinct from how it contributes to processing the outcomes of past choices.

The advent of optogenetics affords the ability to test this hypothesis by examining BLA involvement in decision making during distinct components of the decision process. Hence, the experiments herein examined the effects of BLA inhibition during the deliberation and outcome phases of a risky decision making task involving risk of explicit punishment.

## Materials and Methods

### Subjects

Male Long-Evans rats (weighing 250-275 g upon arrival; Charles River Laboratories, Raleigh, NC) were individually housed and kept on a 12 h light/dark cycle with free access to food and water except as indicated below. Upon arrival, rats were handled daily for one week prior to undergoing surgery. During behavioral testing, rats were maintained at 90% of their free-feeding weight, with their target weights adjusted upward by 5 g/week to account for growth. Animal procedures were conducted in accordance with the University of Florida Institutional Animal Care and Use Committee and followed guidelines of the National Institutes of Health.

### Apparatus

Behavioral testing was conducted in three computer-controlled operant test chambers (Coulbourn Instruments), each of which was contained in a sound-attenuating cabinet. Chambers were equipped with a centrally located food trough (TAMIC Instruments) that projected 3 cm into the chamber and contained a photobeam to detect trough entries. The trough was connected to a feeder, from which 45 mg food pellets (Test Diet, AIN-76A, 5TUL) were delivered into the trough. A nosepoke hole was located above the food trough and two retractable levers were positioned to the left and right of the trough, 11 cm above the floor of the chamber. A 1.12 W lamp was positioned on the back wall of the sound-attenuating cabinet, and served as a houselight. The floor of the test chamber was comprised of stainless steel rods connected to a shock generator that delivered scrambled footshocks. Each operant test chamber was interfaced with a computer running Graphic State 4.0 software (Coulbourn Instruments), which controlled chamber hardware (e.g., lever insertion, nosepoke illumination, food pellet delivery) and recorded task events.

### Laser stimulation

During behavioral test sessions, laser light (560 nm, 8-10 mW output, Shanghai Laser & Optics Century Co., Ltd.) was delivered bilaterally into the BLA in rats expressing halorhodopsin (eNpHR3.0 group) or mCherry (control group) in the BLA. To reach the brain, light was passed from the laser through a patch cord (200 μm core, Thor Labs), a rotary joint (1 X 2, 200 μm core, Doric Lenses) located above the operant chamber, 2 additional patch cords (200 μm core, 0.22 NA, Thor Labs) and bilateral optic fibers (200 μm core, 0.22 NA, 8.3 mm in length; Precision Fiber Products) implanted in the BLA. The laser was interfaced with the computer running Graphic State 4.0 software to allow for precise timing of light delivery during different task phases.

### Surgical procedures

Rats were anesthetized with isoflurane gas (1-5% in O_2_) and received subcutaneous injections of meloxicam (2 mg/kg), buprenorphine (0.05 mg/kg), and sterile saline (10 mL). Rats were placed into a stereotaxic apparatus (David Kopf) and the scalp was cleaned with a chlorohexidine/isopropyl alcohol swab. A sterile adhesive surgical drape was subsequently placed over the body.

For rats used in *in vitro* electrophysiology experiments, the scalp was incised and retracted and the skull was leveled to ensure that bregma and lambda were in the same horizontal plane. Two burr holes were drilled for bilateral virus injections into the BLA (AP: −3.2, ML: ±4.9, DV: −8.5, −8.1 mm from skull surface). At each site, an injection needle was lowered to the target depth and AAV5-CAMKIIα-eHpNR3.0-mCherry (University of North Carolina Vector Core) was infused into the BLA (0.4 μl at the ventral DV coordinate and 0.2 μl at the dorsal DV coordinate, at a rate of 0.5 μl/min). The injection needle was attached to polyethylene tubing, which was connected to a 10 μl Hamilton syringe mounted on a syringe pump (Harvard Apparatus). After each injection, the needle was left in place for an additional 5 minutes to allow for diffusion of the virus. The incision was then sutured and rats were given an additional 10 mL of saline before being placed on a heating pad to recover from surgery.

For rats used in behavioral experiments, the scalp was incised and retracted and six small burr holes were drilled into the skull for placement of jeweler’s screws. Two screws were placed anterior to bregma, two between bregma and lambda and two posterior to lambda. This configuration was used to ensure that the headcap was secured evenly across the skull surface. After leveling the skull to ensure that bregma and lambda were in the same horizontal plane, two additional burr holes were drilled for bilateral implantation of guide cannulae (22 gauge; Plastics One) above the BLA (AP: −3.3, ML: ±4.9, DV: −7.3 from skull surface). Dental cement was used to anchor the cannulae in place. Once the dental cement was set, an injection needle was lowered into each cannula (the tip of the injection needle extended 1.5 mm beyond the end of the cannula) and AAV5-CAMKIIα-eHpNR3.0-mCherry or AAV5-CAMKIIα-mCherry (University of North Carolina Vector Core) was infused into the BLA (0.6 μl at a rate of 0.5 μl/min). A sterile stylet was inserted into each cannula at the completion of the injections. Rats were given an additional 10 mL of saline and were placed on a heating pad to recover from surgery. Rats were allowed to recover for one week before being food restricted in preparation for behavioral testing.

### In vitro electrophysiology

Rats (n = 4) were anesthetized with an intraperitoneal injection of a 75-100 mg/kg ketamine and 5-10 mg/kg xylazine solution and were decapitated using a small animal guillotine. Their brains were rapidly extracted and coronal sections containing the BLA (300 μ m thick) were obtained using a Leica VT 1000s vibratome while submerged in ice cold sucrose laden oxygenated artificial cerebrospinal fluid (aCSF) containing in (mM): 2 KCl, 1.25 NaH2PO4, 1 MgSO_4_, 10 D-glucose, 1 CaCl_2_, 206 sucrose, 25 NaHCO_3_. Slices were then incubated for 30 minutes at 37°C in aCSF which contained in (mM): 124 NaCl, 2.5 KCl, 1.23 NaH_2_PO_4_, 3 MgSO_4_, 10 D-glucose, 1 CaCl_2_, and 25 NaHCO_3_. Following this incubation period slices were allowed to equilibrate to room temperature for a minimum of 30 minutes prior to being used for experiments. All solutions were saturated with 95 % O_2_/ 5 % CO_2_ to maintain a pH of 7.3. For whole cell patch clamp recordings, slices were transferred to a slice chamber where they were continuously perfused at a rate of 1.5-2ml/min with an aCSF bath solution that contained (in mM): 126 NaCl, 3 KCl, 1.2 NaH_2_PO_4_, 1.5 MgSO_4_, 11 D-glucose, 2.4 CaCl_2_ and 25 NaHCO_3_. This solution was also saturated with 95 % O_2_/ 5 % CO_2_ to maintain a pH of 7.3 and bath temperature was maintained at 30-32°C. Slices were visualized using infrared differential interference contrast (IR-DIC) microscopy with an Olympus BX51WI upright stereomicroscope, a 12-bit IRC CCD camera (QICAM Fast 1394, QImaging), and a 40x water immersion lens. Patch pipettes were prepared with a Flaming/Brown type pipette puller (Sutter Instrument, P-97) from 1.5 mm/0.8 mm borosilicate glass capillaries (Sutter Instrument) and pulled to a tip resistance of 4-7 MΩ. Whole cell patch clamp recordings were performed using an Axon Mutliclamp 700B amplifier (Molecular Devices, Sunnyvale, CA) and data were collected at 20 kHz, filtered at 2 KHz and recorded with a Digidata 1322A using Clampex v. 9 or 10 (Molecular Devices, Sunnyvale, CA). BLA neurons expressing mCherry were identified using an epifluoresence microscopy XF102-2 filter set (Omega Optical, excitation: 540-580 nm, emission: 615-695 nm). The light source for epifluoresence microscopy was an X-Cite Series 120Q (Lumen Dynamics). Whole cell patch clamping was initiated under IR-DIC using a potassium-based internal solution that contained (in mM): 130 K-gluconate, 10 KCl, 5 NaCl, 2 MgCl2, 0.1 EGTA, 2 Na2-ATP, 0.3 NaGTP, 10 HEPES and 10 phosphocreatine, pH adjusted to 7.3 using KOH and volume adjusted to 285–300mOsm. Halorhodopsin was activated using 1000 msec light pulses, delivered through the excitation filter in the XF102-2 filter set. Experiments were performed in voltage clamp (at −70 mV), in current clamp (at I=0), or in current clamp during 100-200 pA current injection that was sufficient to drive action potentials. Data were analyzed using custom software written in OriginC (OriginLab, Northampton, MA) by CJF.

### Behavioral procedures

#### Risky Decision-Making Task

Rats were initially shaped to perform the various components of the decision-making task (e.g., lever pressing; nosepoking to initiate a trial) as described previously (Orsini et al., 2015a). They then began training in the Risky Decision-Making task (RDT), which was comprised of three 28-trial blocks and lasted 56 min in duration [this task design was a modification of a similar design used in our laboratory (Simon et al., 2009; Orsini et al., 2015a)]. Each 40 s trial (Figure 1A) began with illumination of the nosepoke and houselight. Upon nosepoking, the nosepoke light was extinguished and either a single lever (forced choice trials) or both levers (free choice trials) extended into the chamber. If rats failed to nosepoke within 10 s, the trial was considered an omission. A press on one lever (left or right; counterbalanced across rats) always yielded a small, “safe” food reward (one food pellet) and a press on the other lever always yielded a large, “risky” food reward (2 food pellets). Delivery of the large reward was accompanied by a variable probability of punishment in the form of a mild footshock (0.25-0.6 mA). The probability of punishment was contingent on a preset probability specific to each block of trials: the probability in the first block was set to 0% and increased across successive blocks (25%, 75%, respectively). The large food reward was delivered irrespective of punishment delivery. Although the levers were counterbalanced across rats, the identities of the small, “safe” lever and large, “risky” lever remained constant for each rat throughout testing. Each block of trials started with eight forced choice trials in which a single lever was extended into the chamber. It is through these forced choice trials that the punishment contingencies for that block were established (four presentations of each lever, randomly presented). During forced choice trials, the probability of punishment following a press for the large reward was dependent upon the outcomes of the other forced choice trial lever presses in that block. For example, in the 25% block, one and only one of the four forced choice trials (randomly selected) resulted in footshock. Similarly, in the 75% block, three and only three of those forced choice trials resulted in footshock. The forced choice trials were followed by 20 free choice trials in which both levers were extended. If rats failed to lever press within 10 s, the house light was extinguished and the trial was counted as an omission. In contrast to the forced choice trials, the probability of punishment in free choice trials was independent, such that the shock probability on each trial was the same regardless of shock delivery on previous trials in that block. During RDT training, shock intensities were adjusted individually for each rat to ensure that there was sufficient parametric space to observe either increases or decreases in risk taking during optogenetic inhibition of BLA.

**Figure 1.**
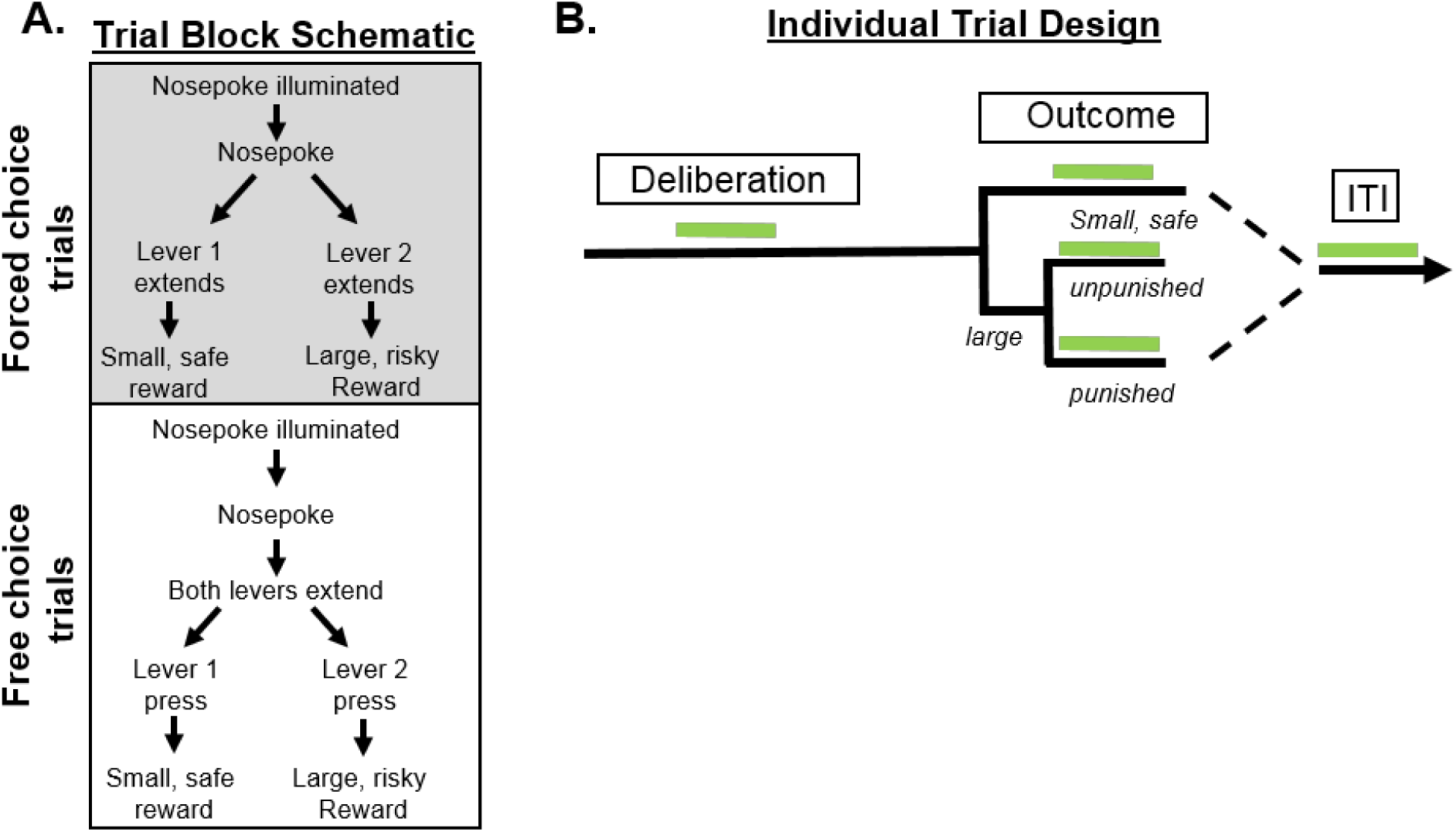
Design of the Risky Decision-Making Task. **A.** Each block consists of 8 forced choice trials and 20 free choice trials. Each free choice trial consists of a deliberation and outcome phase. Rats must nosepoke for the extension of either one lever (forced choice trial) or both levers (free choice trial). A press on one lever yields a small safe reward and a press on the other yields a large reward accompanied by variable probabilities of footshock punishment. **B.** Optogenetic stimulation occurred at one of five possible phases during each free choice trial (with stimulation during each test session taking place in only one of the five phases). Green bars indicate periods of laser stimulation.

Upon reaching stable baseline performance (see *Experimental design and statistical analysis* section for description of stability), rats were lightly anesthetized and optic fibers were inserted into the BLA cannulae such that they extended 1 mm beyond the tips of the cannulae. The fibers were cemented into position and dust caps were placed on the fibers to keep them free from debris. In each subsequent RDT session, spring-insulated patch cords fastened to the rotary joint were attached to the implanted fibers in the rat. Rats were trained in this manner until their performance returned to baseline levels (approximately 3 sessions). Upon reaching this criterion, optogenetic manipulations during test sessions began (note that shock intensities were not adjusted between baseline and laser stimulation sessions). Laser stimulation occurred during three different free choice trial phases (Figure 1B): 1) deliberation 2) reward outcome and 3) intertrial interval (ITI). The deliberation phase consisted of the time between the nosepoke to trigger lever extension and a lever press, and thus captured the period in which rats were presumably deciding between the two available options. Laser stimulation commenced 0.5 s prior to nosepoke illumination and remained on until a lever press occurred or 5 s elapsed, whichever occurred first. For the reward outcome phase, there were three different stimulation conditions: 1) delivery of the small safe reward 2) delivery of the large reward without punishment and 3) delivery of the large reward with punishment. During each outcome condition, laser stimulation began as soon as the rat pressed the lever to yield that outcome and lasted for 5 s. Finally, during the ITI phase, laser stimulation (5 s) occurred 8-15 s after each reward delivery. A randomized, within-subjects design was used such that each rat was tested across multiple stimulation phases. Because of attrition due to detachment of headcaps, however, not all rats were tested for all phases. In between each stimulation session, rats were tethered and tested in the RDT until their performance in the task across two consecutive sessions was no different from their original baseline prior to any stimulation. If choice performance shifted during these re-baselining sessions, shock intensities were adjusted until performance was comparable to the original baseline.

#### Determination of Shock Intensity Threshold

Upon completion of testing in the RDT, rats in the eNpHR3.0 group underwent test sessions in which their shock reactivity was assessed under stimulation and non-stimulation conditions. The procedures were based on those developed by Bonnet and Peterson (1975) to determine the shock thresholds at which specific motor responses were elicited. These test sessions occurred across two days, with each day consisting of two tests: one with laser stimulation and the other without laser stimulation. The order of the test sessions on each day was counterbalanced across the two days. Irrespective of stimulation condition, each test session began with a 2 min baseline period followed by delivery of an unsignaled footshock (0.4 mA, 1 s), which decreased spontaneous motor activity and facilitated detection of motor responses at subsequent low shock intensities. The shock intensity was then set to 0.05 mA and a series of five footshocks (1 s each), each separated by 10 s, was delivered. After each series of footshocks, the shock intensity was increased by 0.025 mA. The increase in shock intensities continued until all motor responses of interest were observed. The shock intensity threshold for a given motor response was determined by the shock intensity at which the given response was elicited by three out of the five footshocks in a series. The motor responses for which shock thresholds were determined consisted of 1) flinch of a paw or a startle response 2) elevation of one or two paws 3) rapid movement of three or all paws. For test sessions with laser stimulation, light was delivered bilaterally (560 nm, 8-10mW) using the same procedures and system used during decision-making sessions. To mimic parameters used for laser stimulation during delivery of the large, punished outcome, laser stimulation and footshock were delivered concomitantly, but the laser remained on for an additional 4 s (total stimulation time of 5 s). Even though no light was delivered during test sessions without laser stimulation, rats were still tethered for the duration of the test.

### Histology and immunohistochemistry

Upon completion of behavioral testing, rats were overdosed with Euthasol and transcardially perfused with cold 0.1M phosphate-buffered saline (PBS) followed by cold 4% paraformaldehyde. Brains were extracted and post-fixed in 4% paraformaldehyde for 24 h before being transferred into a 20% sucrose in 0.1M PBS solution. Brains were sectioned on a cryostat (35 μm) maintained at −20°C. Coronal sections (30 μm) were collected in a 1-in-4 series and placed in wells filled with 0.1M PBS.

Immunohistochemistry was performed on free-floating tissue sections and began with three 10 min washes in 0.1M Tris-buffered saline (TBS). Tissue was then incubated in 3% normal donkey serum (NDS) and 0.3% Triton-X-100 in 0.1M TBS for 1 h at room temperature. Tissue was then immediately transferred into primary antibody [rabbit anti-mCherry at 1:1000 (ab167453, Abcam solution) in 3% NDS and 0.3% Triton-X-100] for 72 hours at 4°C. After primary antibody incubation, tissue was washed three times in 0.1M TBS for 10 min and then incubated in secondary antibody solution [donkey anti-rabbit conjugated to Alexa Fluor 488 at 1:300 (A-21206, Invitrogen) in 3% NDS and 0.3% Triton-X-100) for 2 h at room temperature. Finally, tissue was washed three times in 0.1M TBS for 10 min and then mounted onto electrostatic slides (Fisherbrand) in 0.1M TBS. Slides were coverslipped with Prolong Gold Antifade Mountant (P36941, Invitrogen) and sealed with clear nail polish.

### Experimental design and statistical analyses

Using pilot data collected from several eNpHR3.0 rats, a power analysis was conducted with G*Power software. This analysis indicated that a sample size of at least 4 rats was required to detect significant differences between baseline and stimulation conditions with effect sizes of 0.8 and above, assuming an alpha level of 0.05. To account for possible attrition over the course of the experiment, group sizes were larger than that calculated from the power analysis. A total of 35 male Long-Evans rats were used in these experiments. Twenty-six rats received intra-BLA microinjections of the viral vector containing eNpHR3.0, four of which were used for *in vitro* electrophysiology experiments. Nine rats received intra-BLA microinjections of the viral vector containing mCherry. Within the eNpHR3.0 group, some rats did not undergo every stimulation session due to illness or detachment of headcaps over the course of the experiment. In addition, only a subset of rats (n=6) was used for shock threshold testing. In the control group, there was attrition due to illness or detachment of headcaps, resulting in only four of the initial nine rats completing the stimulation sessions. All 4 rats, however, completed all stimulation conditions.

Raw data files were analyzed using a customized analysis template written in Graphic State 4.0 software. This template extracted data for specific task events of interest: numbers of lever presses during forced and free choice trials, latencies to press levers, latencies to nosepoke, and numbers of omissions during forced and free choice trials. The behavioral and statistical procedures were conducted identically for the eNpHR3.0 and control groups. Choice performance in each block of the RDT was measured as the percentage of free choice trials (each block consisted of 20 free choice trials; excluding omissions) on which rats chose the large, risky outcome. Each rat was trained in the RDT until it reached stable baseline performance. Stable baseline was obtained when the coefficient of variation (CV) for choice of the large, risky outcome was less than 20% in each block for at least two consecutive sessions. Once this criterion was met, stimulation sessions commenced. In between each stimulation session, rats were re-trained in the RDT until their behavior re-stabilized, which was determined using the same criterion. To ensure that the baseline after stimulation was similar to the original baseline (before any stimulation sessions took place), the CV of the means of each block between baseline sessions had to fall below 20%. Upon reaching this criterion, rats were advanced to the next stimulation session. Effects of stimulation (i.e., BLA inhibition) on choice performance were determined using a two-factor repeated measures ANOVA with session condition (i.e., baseline vs. inhibition) and trial block as within-subjects factors. In all analyses, a *p*-value of 0.05 or less was considered statistically significant. Latencies to nosepoke to trigger lever extension were measured as the interval between the illumination of the nosepoke light and a nosepoke response, excluding trials on which the rat failed to nosepoke altogether (omissions). Using a repeated measures ANOVA, nosepoke response latencies were specifically compared between baseline and deliberation stimulation sessions to determine whether laser stimulation (which was initiated 0.5 sec before nosepoke illumination) affected this aspect of behavior. Effects of BLA inhibition on omissions during free and forced choice trials were analyzed using a paired *t*-test with session condition as the within-subjects factor.

To better understand the effects of BLA inhibition during task phases in which inhibition significantly affected choice behavior, additional analyses were conducted to determine whether optogenetic manipulations altered the degree to which feedback from past trials influenced subsequent choices. Specifically, this analysis provided a measure of how BLA inhibition affected the likelihood of choosing the large, risky outcome upon receipt of the large reward in the absence of punishment on the previous trial (win-stay performance) vs. the likelihood of choosing the large, risky outcome upon receipt of the large reward accompanied by punishment on the previous trial (lose-shift performance; Bari et al., 2011; St Onge et al., 2011). To perform this analysis, choices were categorized according to the outcome of the previous trial (large, punished outcome vs. large, unpunished outcome). Win-stay performance was calculated as the number of trials within each free choice block in which a rat chose the large, risky lever after receipt of a large, unpunished outcome (win), divided by the total number of free choice trials in which the rat received a large, unpunished outcome. Similarly, lose-shift performance was calculated as the number of trials within each free choice block in which a rat chose the small, safe lever after receipt of a large, punished outcome (lose), divided by the total number of free choice trials in which the rat received a large, punished outcome. Effects of BLA inhibition on the percentage of win-stay and lose-shift trials were each analyzed using paired *t-*tests with session (baseline vs. inhibition) as the within-subjects factor.

Shock threshold intensities for the laser stimulation or no laser stimulation sessions were averaged across the two test days. Analysis of shock intensity thresholds was conducted using a two-factor repeated measures ANOVA with stimulation condition (inhibition vs. no inhibition) and motor response as the within-subjects factors. To eliminate the possibility that the order of the test sessions on each day contributed to differences in shock reactivity thresholds, another repeated measures ANOVA was conducted using the same within-subjects factors and also included order of laser stimulation as a between-subjects factor. If either of these parent ANOVAs resulted in main effects or significant interactions, additional repeated measures ANOVA or paired *t*-tests were performed to determine the source of significance.

## Results

### In vitro electrophysiology

In slices from rats injected with AAV5-CAMKIIα-eHpNR3.0-mCherry, BLA neurons expressing mCherry were identified with epifluorescence microscopy and recorded from using conventional whole-cell recording techniques (see *Methods* section). mCherry-positive BLA neurons (n=11) had a mean whole cell capacitance of 149 ± 14.9 pF. A subset of these neurons was filled with biocytin, immunolabeled with Alexa-594, and imaged with 2-photon meditated epifluorescence microscopy. Cells examined in this manner were all multipolar and had dense local dendritic branches within the BLA (Figure 2A). Collectively, these features are consistent with effective transduction of glutamatergic BLA principal neurons. A 1 s activation of eHpNR3.0 in mCherry-positive BLA neurons voltage clamped at −70 mV (see *Methods* section) produced a clear outward current which had a peak amplitude of 117 ± 29.6 pA, obtained within ∼100 msec of activation, and a mean amplitude of 80.0 ± 20.8 pA as observed during the last 200 msec of activation (Figure 2B). Identical stimulation in current clamp (I=0) produced a maximum hyperpolarization of −16 ± 3.1 mV (also obtained within ∼100 msec of activation), and a mean hyperpolarization of −9.0 ± 2.0 mV as observed during the last 200 msec of activation (Figure 2C). This hyperpolarization was sufficient to completely silence 9 out of 11 cells tested when firing under a 100-200 pA load (Fig. 1D). Firing rate was slowed, but not eliminated, in the other two cells. Collectively, these results demonstrate that activation of eHpNR3.0 produces robust functional inhibition of BLA principal neurons.

**Figure 2.**
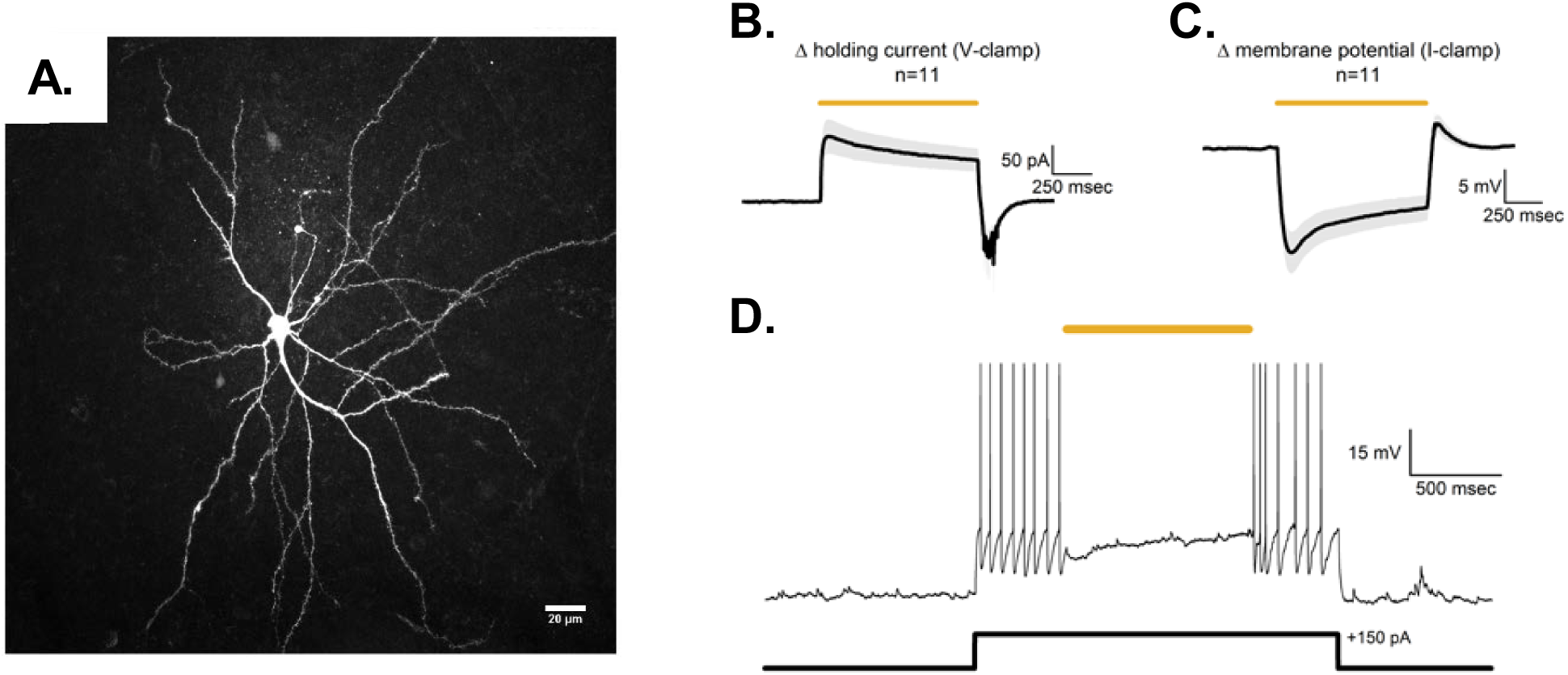
Functional validation of eNpHR3.0 in the BLA. **A.** Two-photon z-series projection of a mCherry-positive BLA neuron filled with biocytin and immunolabeled with Alexa-594. **B.** mCherry-positive cells (n=11) voltage clamped at −70 mV show an increase in holding current upon light stimulation (yellow line) and return to baseline holding current upon light termination. Black line indicates the mean response and shaded area indicates SEM. **C.** mCherry-positive cells (n=11) current clamped at 0 pA show hyperpolarization upon light stimulation and return to resting membrane potential upon light termination. Black line indicates the mean response and shaded area indicates SEM. **D.** A representative mCherry-positive cell that was current clamped at 0 pA shows an increase in firing rate upon injection of a +150 pA current pulse, which is effectively suppressed during light delivery.

### Histology

Of the 22 rats that received the viral vector containing eNpHR3.0 for optogenetic manipulations, one died during surgery and five were euthanized during training due to detached headcaps. Of the remaining 16 rats, three were excluded due to off-target fiber placements (too ventral; n = 1) or lack of eNpHR3.0 expression in one hemisphere (n = 2). Figure 3A displays the maximum (light gray) and minimum (dark gray) spread of the virus, and Figure 3B depicts the location of optic fiber tips of rats that were included in the final data analysis. A representative placement of a fiber tip in the BLA with eNpHR3.0 expression is shown in Figure 3C.

**Figure 3.**
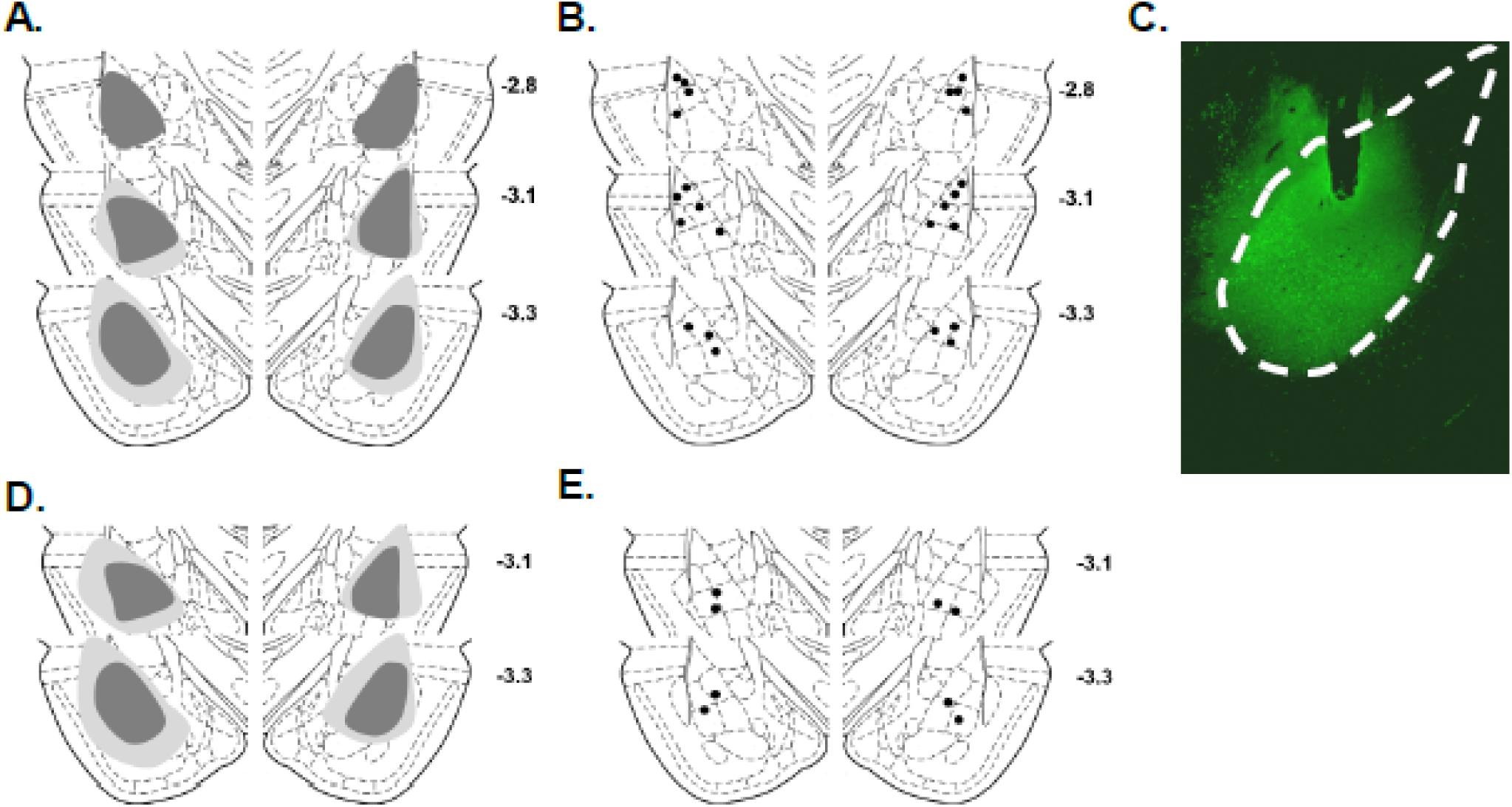
eNpHR3.0 expression and optic fiber placement in the BLA. **A.** Schematic depicting the maximum (light gray) and minimum (dark gray) spread of eNpHR3.0 expression in the BLA. **B.** Optic fiber placements in the BLA. Black circles represent the tips of the optic fibers. **C.** Representative micrograph depicting eNpHR3.0 expression and the tip of the optic fiber in the BLA. Dashed white line represents the borders of the BLA. **D.** Schematic depicting the maximum (light gray) and minimum (dark gray) spread of mCherry expression in the BLA of control rats. **E.** Optic fiber placements in the BLA in control rats. Black circles represent the tips of the optic fibers.

Of the 9 rats that received the viral vector containing mCherry alone, one died during surgery and four were euthanized during training due to detached headcaps, resulting in a final n = 4. Figure 3D displays the maximum (light gray) and minimum (dark gray) spread of the virus and Figure 3E shows the location of optic fiber tips of control rats that were included in the final data analysis.

### Optogenetic BLA inhibition during decision making in eNpHR3.0 rats

#### BLA inhibition during deliberation

Optogenetic inhibition of the BLA during deliberation (n = 12) caused a significant decrease in choice of the large, risky outcome [decreased risky choice; inhibition, *F* (1, 11) = 14.57, *p* < 0.01; inhibition X trial interaction [*F* (2, 22) = 10.29, *p* < 0.01; Figure 4A]. Importantly, this effect was only observed in blocks of trials in which there was a risk of punishment: while there was no effect of inhibition in block 1 [*t* (11) = −1.27, *p* = 0.23], BLA inhibition decreased choice of the large, risky outcome in both block 2 [*t* (11) = 4.51, *p* < 0.01] and block 3 [*t* (11) = 2.16, *p* = 0.05].

**Figure 4.**
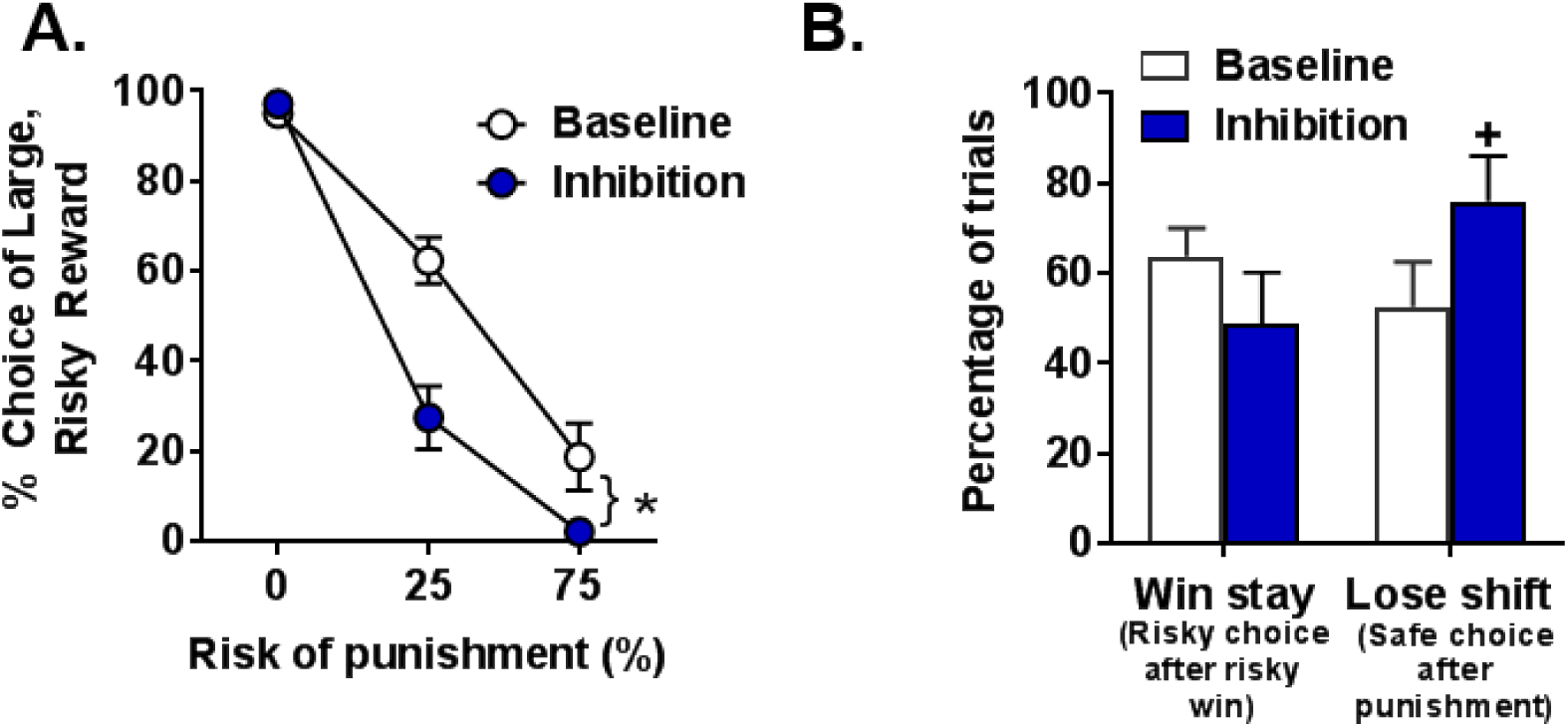
BLA inhibition during deliberation decreases risky choice. **A.** BLA inhibition decreases choice of the large, risky outcome. **B.** There were no effects of BLA inhibition on winstay trials. In contrast, there was a near-significant increase in lose-shift trials upon BLA inhibition. Data are represented as mean ± standard error of mean (SEM). An asterisk indicates a significant difference and a cross indicates a trend (*p* = 0.08) toward a significant difference between inhibition and baseline (no laser) conditions.

Additional analyses were performed to determine whether BLA inhibition during deliberation affected the percentage of win-stay or lose-shift trials (Figure 4B). There was no effect of inhibition on the percentage of win-stay trials (*t* (8) = 1.61, *p* = 0.15), but there was a near significant increase in the percentage of lose-shift trials (*t* (9) = −1.99, *p* = 0.08). Note that in the win-stay analysis, three rats were excluded because they either never chose the large, risky outcome or never encountered a trial in which they chose the large, risky outcome and received the large reward without punishment. Similarly, in the lose-shift analysis, two rats were excluded because they never selected the large, risky outcome. This slight increase in lose-shift trials suggests that BLA inhibition slightly increased the likelihood for rats to shift their choice to the small, safe outcome after receiving a large reward accompanied by punishment. Collectively, these results show that BLA inhibition during the period in which rats deliberated between the two available options caused an increase in risk aversion.

Finally, there was no effect of BLA inhibition during deliberation on omissions in either the forced choice trials [*t* (11) = −0.87, *p* = 0.44] or the free choice trials [*t* (11) = −0.26, *p* = 0.80]. Additional analyses were conducted to determine whether inhibition affected rats’ latency to nosepoke to trigger lever extension. While there was no main effect of inhibition [*F* (1, 11) = 0.05, *p* = 0.82], there was a trend toward a significant inhibition X block interaction [*F* (2, 22) = 3.10, *p* = 0.07], with BLA inhibition causing a slight decrease in latency to nosepoke, particularly in block 3 [mean of 1.87 (±0.24) s for baseline; mean of 1.48 (± 0.15) s for stimulation]. Note, however, that because light onset commenced 0.5 s before the nosepoke was illuminated to signal the beginning of a trial, BLA inhibition should have been maximal prior to the start of the deliberation period.

#### BLA inhibition during delivery of the small, safe outcome

Optogenetic inhibition of the BLA during delivery of the small, safe outcome (n = 10) had no effect on choice of the large, risky outcome [inhibition, *F* (1, 9) = 0.09, *p* = 0.77; inhibition X block, *F* (2, 18) = 1.73, *p* = 0.21; Figure 5A]. Additionally, inhibition had no effect on omissions (Table 1) during forced choice trials [*t* (9) = −0.32, *p* = 0.76] or during free choice trials [*t* (9) = −0.91, *p* = 0.39]. Hence, BLA inhibition during delivery of the small, safe outcome did not alter choice behavior.

**Figure 5.**
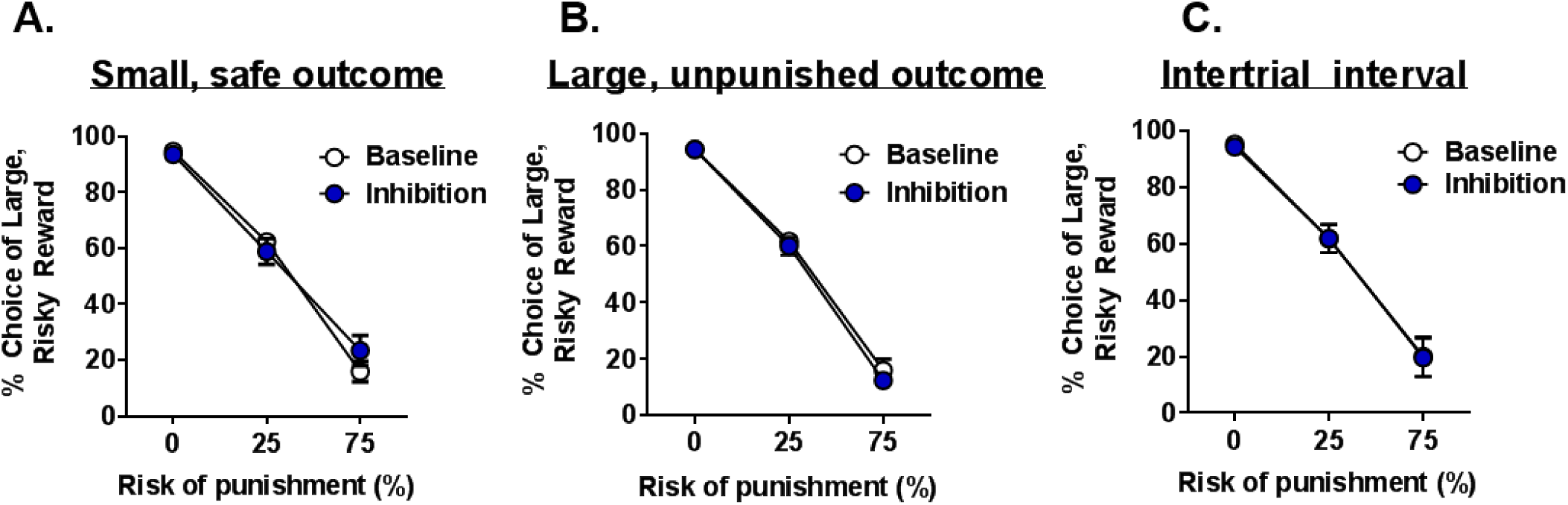
BLA inhibition has no effect on risky choice during other task phases. **A.** BLA inhibition during delivery of the small, safe outcome did not affect choice of the large, risky outcome. **B.** BLA inhibition during delivery of the large, unpunished outcome had no effect on choice of the large, risky outcome. C. BLA inhibition during the ITI had no effect on choice of the large, risky outcome. Data are represented as mean ± SEM.

#### BLA inhibition during delivery of the large, unpunished outcome

Similarly, there was no effect of BLA inhibition during the large, unpunished outcome (n = 9) on choice behavior [inhibition, *F* (1, 8) = 0.45, *p* = 0.52; inhibition X block, *F* (2, 16) = 0.30, *p* =0.74; Figure 5B]. There were also no effects of inhibition on omissions (Table 1) during forced choice trials [*t* (8) = 0.50, *p* =0.63] or free choice trials [*t* (8) = −1.0, *p* = 0.35]. Collectively, these findings indicate that BLA inhibition during delivery of the large, unpunished outcome did not affect choice behavior.

#### BLA inhibition during delivery of the large, punished outcome

In contrast to BLA inhibition during delivery of the large, unpunished outcome, optogenetic BLA inhibition during delivery of the large, punished outcome (n = 10) significantly increased choice of the large, risky outcome [inhibition, *F* (1, 9) = 82.75, *p* < 0.01; inhibition X block, *F* (1, 9) = 39.22, *p* < 0.01; Figure 6A]. It is important to note that this analysis only used choice behavior in the 25% and 75% blocks from baseline and stimulation sessions, as they were the only blocks in which BLA inhibition could occur.

**Figure 6.**
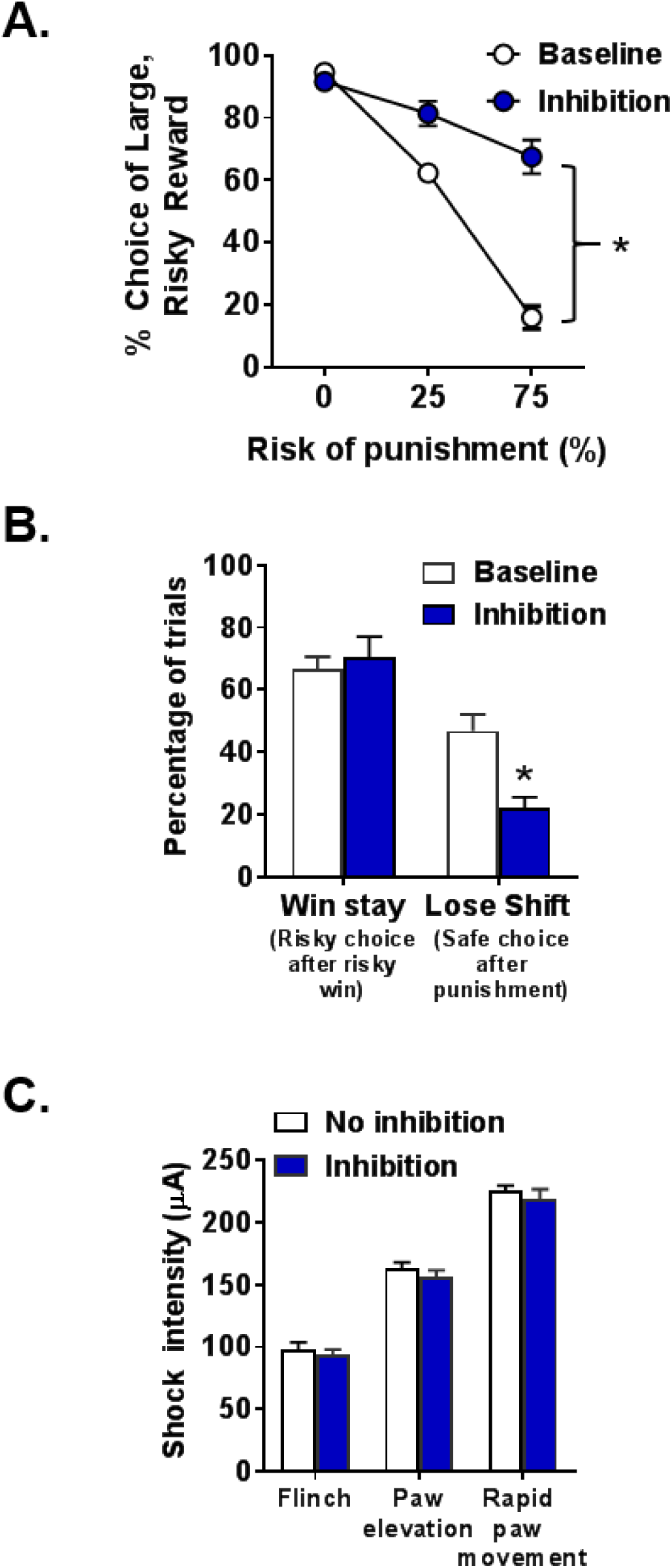
BLA inhibition during delivery of the large, punished outcome increases risky choice. **A.** BLA inhibition increased choice of the large, risky outcome. **B.** There was no effect of BLA inhibition on win-stay performance. In contrast, BLA inhibition decreased lose-shift performance. **C.** BLA inhibition did not alter the intensity thresholds at which shock elicited a flinch, elevation of 1-2 paws, or rapid movement of 3 or all paws. Data are represented as mean ± SEM. Asterisks indicate a significant difference between inhibition and baseline (no laser) conditions.

Given the significant effects of BLA inhibition during this phase of the task, additional analyses were performed to determine how this manipulation affected the percentage of win-stay or lose-shift trials (Figure 6B). There was no effect of BLA inhibition on the percentage of win/stay trials [*t* (9) = −0.44, *p* = 0.67]; however, there was a significant decrease in the percentage of lose/shift trials [*t* (9) = 3.02, *p* = 0.01] compared to baseline. Thus, BLA inhibition during delivery of the large, punished outcome caused rats to increase the likelihood of choosing the large, risky outcome, despite having been punished for this choice on the preceding trial.

Lastly, there were no effects of BLA inhibition on omissions (Table 1) during free choice trials [*t* (9) = 0.09, *p* = 0.93], although inhibition did cause a significant decrease in omissions during forced choice trials compared to baseline conditions [*t* (9) = 2.56, *p* = 0.03].

#### BLA inhibition during shock threshold testing

Rather than affecting processes related to risk taking *per se*, the effects of BLA inhibition during delivery of the large, punished outcome may have been due to an inhibition-induced decrease in shock sensitivity. To address this, a subset of rats (n = 6) was tested in a behavioral assay that evaluates the thresholds at which selective motor responses (as described in the *Methods* section) are elicited by shock delivery. These thresholds were obtained under stimulation and no stimulation (inhibition vs. no inhibition, respectively) conditions (Figure 6C). A two-factor repeated measures ANOVA revealed neither a main effect of inhibition [*F* (1, 5) = 4.00, *p* = 0.10] nor an inhibition X motor response interaction [*F* (2, 10) = 0.04, *p* = 0.96]. Thus, the increase in risky choice during sessions in which BLA inhibition occurred during delivery of the large, punished outcome cannot be accounted for by a decrease in footshock sensitivity.

#### BLA inhibition during ITIs

Optogenetic inhibition of the BLA during the ITI (n = 13) had no effect on choice of the large, risky outcome [inhibition, *F* (1, 12) = 0.01, *p* = 0.91; inhibition X block, *F* (2, 24) = 0.02, *p* = 0.98; Figure 5C]. Similarly, BLA inhibition during ITIs did not affect omissions during forced choice trials [*t* (12) = 0.3, *p* = 0.77], but caused a near significant increase in omissions during free choice trials [*t* (12) = −2.04, *p* = 0.06].

### Optogenetic BLA stimulation during decision making in control rats

To ensure that the effects of BLA inhibition were not due to light delivery alone, another group of rats received intra-BLA microinjections of a vector carrying mCherry alone and were then trained in the RDT. Because BLA inhibition only altered choice behavior during deliberation and delivery of the large, punished outcome in eNpHR3.0 rats, control rats only received stimulation during these two phases (in separate sessions, in a randomized order across rats).

#### BLA stimulation during deliberation

BLA stimulation during deliberation (n = 4) had no effect on choice of the large, risky outcome compared to baseline conditions [stimulation, *F* (1, 3) = 1.00, *p* = 0.39; stimulation X block, *F* (2, 6) = 1.00, *p* = 0.42; Figure 7A]. There was no main effect of BLA stimulation on latency to nosepoke to initiate lever extension [*F* (1, 3) = 2.33, *p* = 0.23; Table 1]; however, it appeared that under stimulation conditions, latency to nosepoke did increase across the session [*F* (2, 6) = 7.70, *p* = 0.02]. While there was a trend toward a significant effect of stimulation on omissions (Table 1) during forced choice trials [*t* (3) = −2.82, *p* = 0.07], this was due to the fact that there were fewer omissions under stimulation compared to baseline conditions. There were no effects of BLA stimulation on omissions during free choice trials [*t* (3) = 1.00, *p* = 0.39]. Collectively, these results indicate that laser stimulation of BLA alone during deliberation did not affect risky decision making in control rats.

**Figure 7.**
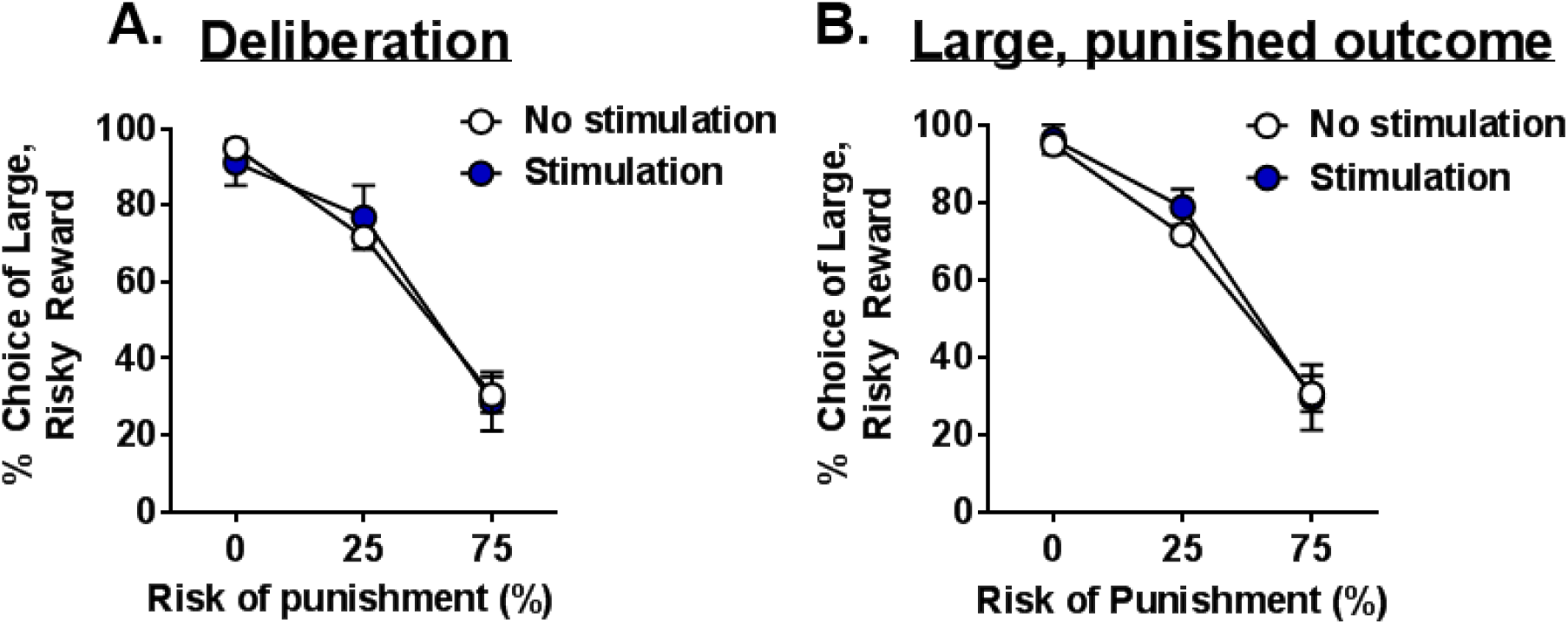
Laser stimulation of the BLA has no effect on risky choice in control rats. **A.** In rats injected with vectors carrying mCherry alone, BLA stimulation during deliberation did not affect choice of the large, risky outcome. **B**. BLA stimulation during delivery of the large, punished outcome had no effect on choice of the large, risky outcome. Data are represented as mean ± SEM.

#### BLA stimulation during delivery of the large, punished outcome

There was also no effect of BLA stimulation during delivery of the large, punished outcome (n = 4) on choice of the large, risky outcome [stimulation, *F* (1, 3) = 1.00, *p* = 0.39; stimulation X block, *F* (2, 6) = 1.0, *p* = 0.42; Figure 7B]. There was a trend toward a significant effect of stimulation on omissions (Table 1) during forced choice trials [*t* (3) = −2.82, *p* = 0.07]; however, this again appeared to be due to fewer omissions under stimulation compared to baseline conditions. There were no differences in omissions during free choice trials between stimulation and baseline conditions [*t* (3) = 1.67, *p* = 0.19]. Hence, laser stimulation of BLA alone during delivery of the large, punished outcome had no effect on risky decision making in control rats.

## Discussion

Decision making involves coordination of multiple cognitive functions to generate choice behavior. While there is a growing body of literature delineating the neural substrates governing decision making, less is known about how and when such brain regions are engaged during the decision process. The current study demonstrates that the BLA plays distinct roles during different components of risky decision making. Whereas optogenetic inhibition of the BLA during deliberation resulted in a decrease in choice of the large, risky outcome (decreased risky choice), BLA inhibition during delivery of the large, punished outcome had the opposite effect (increased risky choice). These effects were specific to the task phase in which inhibition occurred because BLA inhibition had no effect on choice behavior during delivery of the small, safe outcome, the large, unpunished outcome, or the ITI. Further, there were no effects of light delivery into the BLA during deliberation or delivery of the large, punished outcome in control rats (in the absence of eNpHR3.0).

The overall finding that BLA manipulation alters choice performance during risky decision making is consistent with previous studies implicating this region in cost/benefit decision making. In a risky decision making task involving choices between a small, certain food reward and a large, probabilistic food reward, pharmacological inactivation of BLA decreased choice of the large, probabilistic reward, but only at probabilities at which it was more profitable to choose this reward (Ghods-Sharifi et al., 2009). Consistent with this, BLA lesions induce a pattern of disadvantageous choice behavior in another rodent model of risky decision making designed to simulate the Iowa Gambling Task (Zeeb and Winstanley, 2011). More recently, we showed that BLA lesions increase risky choice in the RDT (Orsini et al., 2015a), and control experiments suggested that this increase was due to impaired integration of reward magnitude and punishment-related information. Given the complexity of the decision-making process, however, the use of lesions and pharmacological inactivation, while informative, may obscure a complete understanding of how the BLA is engaged during the course of individual decisions.

To circumvent this issue, the current study employed optogenetics to selectively inhibit the BLA during distinct phases of the decision-making process. In contrast to effects of permanent BLA lesions (Orsini et al. 2015a), optogenetic inhibition caused both an increase *and* decrease in risky choice depending on the timepoint at which inhibition occurred. These results suggest that the contribution of the BLA to risky choice is not uniform, but instead that it may function in different capacities even over the course of a few seconds of a decision-making trial. During deliberation, various sources of information must be assimilated to bias behavior toward a specific choice. In particular, information about the anticipated rewarding aspects of each potential outcome must be integrated and weighed against the negative/adverse aspects of those outcomes. BLA inhibition during this period interfered with this integrative process such that choices were more strongly biased by punishment-related information. One possibility is that this is due to a loss of reward magnitude information, although this seems unlikely given that choice behavior was intact in the first block of trials (in which there was no risk of punishment). Alternatively, and consistent with the slight increase in lose-shift trials, BLA inhibition may have augmented the salience of the punishment associated with the large reward. This also seems unlikely, however, given that lesions and pharmacological inactivation of the BLA reduce fear expression in other contexts (Helmstetter and Bellgowan, 1994; Maren et al., 1996). A final, and more likely possibility is that BLA inhibition during deliberation may have attenuated the incentive salience of anticipated outcomes and, consequently, the ability to bias action selection toward more salient rewards. Hence, the BLA may be important for tagging available outcomes based on their incentive salience (i.e., to favor larger, albeit risker, outcomes). In the absence of an intact BLA, the punishment history and/or aversive properties of these outcomes prevail and drive choice behavior.

In contrast, the increase in risk-taking following BLA inhibition during delivery of the large, punished outcome suggests that the BLA is engaged in a manner different from that during deliberation. Incorporating feedback about outcomes of past choices to guide future choice is a critical aspect of adaptive decision making. The BLA has long been implicated in encoding and representing aversive properties of stimuli in Pavlovian and instrumental learning tasks (Wassum and Izquierdo, 2015). Thus, inhibition during delivery of the large, punished outcome may have prevented the BLA from encoding the punishing aspects of this outcome and therefore impaired the ability to use this information as feedback to adjust future choice behavior. This would result in choice performance being driven by rewarding properties of this outcome, irrespective of whether its delivery was accompanied by footshock. This is supported by the significant decrease in lose-shift trials such that rats continued to choose the large, risky outcome despite having been punished on the preceding trial. Importantly, the effects of BLA inhibition during this phase were not due to alterations in shock sensitivity, as there were no changes in thresholds at which shock-induced motor responses were elicited. This is consistent with previous work showing that BLA lesions do not affect discrimination between punished and unpunished rewards of the same magnitude (Orsini et al., 2015a) and when considered together, demonstrates that the BLA is not necessary for encoding shock alone. Collectively, these data suggest that when a rewarding outcome is accompanied by an adverse consequence, the BLA may be responsible for encoding the negative aspects of that outcome that can then be used as feedback during future deliberation.

The idea that the BLA functions in a heterogeneous manner during risky decision making is consistent with previous work showing that BLA neurons that encode outcomes of different valences are segregated into distinct populations (Schoenbaum et al., 1998; Paton et al., 2006; Belova et al., 2007; Belova et al., 2008; Shabel and Janak, 2009; Sangha et al., 2013; Zhang et al., 2013; Namburi et al., 2015; Beyeler et al., 2016). Aversive and appetitive outcomes are predominantly represented by separate BLA cell populations (Namburi et al., 2015; Beyeler et al., 2016), suggesting that the functional heterogeneity of the BLA during risky decision making could arise from distinct neuronal populations representing incentive salience (positive-value neurons) vs. aversive properties (negative-value neurons) of choice outcomes. The current data further suggest that these separate populations are differentially engaged depending on the phase of the decision process. Thus, positive-value neurons may be important during the deliberative process for signaling the incentive salience of possible outcomes, whereas negative-value neurons may be critical for sensitivity to negative feedback. It is not clear, however, whether these separate populations of neurons interact with one another and if so, when and where this interaction occurs. If, in fact, these distinct neuronal populations are differentially engaged during decision making, how do they ultimately affect choice behavior? One possibility is that the positive- and negative-value neurons have divergent and non-overlapping downstream targets. Indeed, BLA neurons that project to the nucleus accumbens (NAc) selectively support reward conditioning whereas BLA neurons that project to the central nucleus of the amygdala (CeA) selectively support fear conditioning (Namburi et al., 2015; Beyeler et al., 2016). While the BLA-NAc projection is implicated in risky decision making (St Onge et al., 2012), the contribution of the BLA-CeA circuit is unknown. It is also possible that putative positive- and negative-value BLA neurons modulate risky choice through divergent projections to the core and shell subregions of the NAc, respectively. This hypothesis is consistent with the canonical theory that the NAc core (NAcC) is important for facilitating approach behavior whereas the NAc shell (NAcSh) is required for suppressing ongoing behavior (Floresco, 2014). This functional dichotomy extends to instrumental tasks involving conflict or punishment: NAcSh inactivation increases punished responding (Piantadosi et al., 2017) and decreases avoidance responses (Fernando et al., 2014), whereas NAcC inactivation decreases overall reward-seeking, irrespective of accompanying punishment (Piantadosi et al., 2017). Further evidence indicates that these distinct functions are modulated by BLA input. For example, activation of the BLA-NAcC pathway drives reward-seeking behavior (Ambroggi et al., 2008; Stuber et al., 2011; Namburi et al., 2015) and interruption of this circuit impairs reward conditioning and decision making (Ambroggi et al., 2008; Stuber et al., 2011; St Onge et al., 2012). In contrast, the BLA-NAcSh, but not the BLA-NAcC, pathway supports active avoidance behavior (Ramirez et al., 2015). Thus, positive-encoding BLA neurons may contribute to the deliberative process via their downstream connections with the NAcC and negative-encoding BLA neurons may provide negative feedback information through their interactions with the NAcSh. Interestingly, it has been proposed that networks of inhibitory BLA interneurons may play a permissive role in determining which neuronal circuits are engaged during motivated behavior (Janak and Tye, 2015), which could allow flexible shifts in choice behavior as reward or punishment contingencies change.

To our knowledge, this study is the first to demonstrate multiple roles for the BLA in decision making depending on the phase of the decision process engaged. These results highlight the need to use more temporally targeted manipulations to understand the neural circuitry supporting complex cognitive operations. More importantly, these findings provide a more refined understanding of how the BLA contributes to risk-based decision making, and a foundation for future work on development of novel approaches for remediating maladaptive choice behavior.

## Acknowledgements

We thank Mr. Matthew Bruner and Mrs. Shannon Wall for their technical assistance. We also wish to acknowledge the McKnight Brain Research Foundation for their support of this project. The work was further supported by a Thomas H. Maren Fellowship and K99DA041493 (CAO), a McKnight Predoctoral Fellowship and the Pat Tillman Foundation (CMH), and R01DA036534 (BS).

## Notes

**Conflicts of interest**: The authors declare no competing financial interests.

